# Monoacylglycerol O-acyltransferase 1 is required for adipocyte differentiation *in vitro* but does not affect adiposity in mice

**DOI:** 10.1101/2022.02.14.480414

**Authors:** Jason M. Singer, Trevor M. Shew, Daniel Ferguson, M. Katie Renkemeyer, Terri A. Pietka, Angela M. Hall, Brian N. Finck, Andrew J. Lutkewitte

**Affiliations:** Center for Human Nutrition, Washington University School of Medicine, St. Louis, MO

## Abstract

**Objective:** Monoacylglycerol O-acyltransferase 1 (*Mogat1*), a lipogenic enzyme that converts monoacylglycerol to diacylglycerol, is highly expressed in adipocytes and may regulate lipolysis by re-esterifying fatty acids released during times when lipolytic rates are low. However, the role of *Mogat1* in regulating adipocyte fat storage during differentiation and diet-induced obesity is relatively understudied.

**Methods:** Here we generated adipocyte-specific *Mogat1* knockout mice and subjected them to a high-fat diet to determine the effects of *Mogat1* deficiency on diet-induced obesity. We also used *Mogat1* floxed mice to develop preadipocyte cell lines wherein *Mogat1* could be conditionally knocked out to study adipocyte differentiation *in vitro*.

**Results:** In preadipocytes, we found that *Mogat1* knockout at the onset of preadipocyte differentiation prevented the accumulation of glycerolipids and reduced the differentiation capacity of preadipocytes. However, the loss of adipocyte *Mogat1* did not affect weight gain or fat mass induced by high-fat diet in mice. Furthermore, loss of *Mogat1* in adipocytes did not affect plasma lipid or glucose concentrations or insulin tolerance.

**Conclusions:** These data suggest *Mogat1* may play a role in adipocyte differentiation *in vitro* but not adipose tissue expansion in response to nutrient overload in mice.

**STUDY IMPORTANCE:** *What is already known?:*

- Adipose tissue expansion through adipocyte precursor cell differentiation is critical for proper lipid storage during nutrient overload.
- Monoacylglycerol O-acyltransferase 1 (*Mogat1*), a lipogenic enzyme, is highly induced during adipocyte differentiation of human and mouse precursor cells and is reduced in patients with obesity and metabolic dysfunction.

*What does this study add?:*

- *Mogat1* deletion during early adipocyte differentiation reduces differentiation capacity, adipogenic gene expression and lowers glycerolipid content of differentiated adipocytes.
- Adipocyte *Mogat1* expression is dispensable for adiposity and metabolic outcomes high-fat fed mice and suggests compensation from other glycerolipid synthesis enzymes.

*How might these results change the direction of research?:*

- Understanding the molecular mechanisms of glycerolipid metabolism healthy adipose tissue expansion.

## INTRODUCTION

Adipose tissue regulates systemic metabolism through endocrine factors (adipokines) and the regulated storage and release of free fatty acids (FFA) and glycerol. Triglycerides comprise three fatty acids esterified to a glycerol backbone and represent the most abundant lipid species stored within adipocytes. In higher organisms, two pathways have evolved for TAG synthesis that converge at diacylglycerol (DAG) synthesis, the glycerol-3-phosphate and the monoaclylglycerol O-acyltransferase pathways. The monoacylglycerol O-acyltransferase (MGAT) family of enzymes acylate monoacylglycerol (MAG) into DAG; most of which is directly acylated into triglyceride (TAG) via diacylglycerol O-acyltransferase (DGAT) activity. In humans, three genes encode enzymes of this family (*MOGAT1*, *MOGAT2*, and *MOGAT3*), but there are only two *Mogat* family genes (*Mogat1* and *Mogat2*) in mice.

*Mogat2* is enriched in intestinal enterocytes and is important for dietary fat absorption (1–3). In contrast, *Mogat1* is highly induced in liver of mice with hepatic steatosis and is the primary MGAT isoform expressed in adipose tissue of humans and rodents (4–6). While intestinal MGAT function has been studied for many years, less is known about the physiologic function of MGAT in other organs. We have recently shown that hepatic and constitutive *Mogat1* deletion does not affect hepatic steatosis and insulin resistance in mice in the context of diet-induced obesity (7). However, it remains to be determined whether modulating MGAT activity in adipose tissue specifically might have efficacy for preventing or treating obesity-related metabolic disease.

*Mogat1* expression and MGAT activity are highly induced during differentiation of mouse and human precursors cells into mature adipocytes (4). Adipose tissue *Mogat1* gene expression is reduced in genetic mouse models of marked obesity (*db/db* mice) and abdominal fat of people with obesity that exhibit metabolic abnormalities compared to people with metabolically-healthy obesity (4). Expression of *Mogat1* in abdominal adipose tissue also inversely correlates to basal rates of FFA release (a measure of adipose tissue lipolysis) (4). Similarly, mice lacking *Mogat1* in adipose tissue have increased basal lipolysis on standard chow diets (4) and *Mogat1* expression and activity are reduced by fasting when lipolytic rates are high (4). Overall, these data suggest that MGAT activity serves to re-esterify FFA to limit FFA release when lipolytic rates should be low (6, 8) and that the loss of *Mogat1* could lead to “leak” of FFAs into circulation and deposition in other tissues leading to deleterious effects such as insulin resistance in skeletal muscle and liver (9). Indeed, adipose tissue expression of *MOGAT1* positively correlates to glucose disposal rates during a hyperinsulinemic-euglycemic clamp in people with obesity (4). However, this phenomenon is associative only and the cause and effect relationship between these observations requires further study.

Several questions remain regarding the role of *Mogat1* in adipocyte differentiation and function. For example, although *Mogat1* expression is highly induced early in the adipogenic program (4), the requirement for *Mogat1* in adipocyte differentiation remains unclear. Does diminished adipose tissue MGAT activity lead to ectopic fat accumulation and metabolic abnormalities with nutrient overload? Few studies have addressed the requirement for *Mogat1* in adipose tissue expansion during the progression of obesity (10–12) and those that have, used global knockout of *Mogat1* rather than deleting in an adipocyte-specific manner. We have recently shown global deletion of *Mogat1* actually leads to unexpected increases in weight gain in mice fed a high-fat diet (7). Herein, we sought to determine the functional consequences of *Mogat1* depletion during adipocyte differentiation *in vitro* and expansion *in vivo* by using mice harboring a floxed allele of *Mogat1* and by generating adipocytes with conditional deletion of *Mogat1*.

## METHODS

### Animal studies

All mouse studies were approved by the Institutional Animal Care and Use Committee of Washington University. *Mogat1* whole body null mice and adipocyte-specific *Mogat1* knockout mice were generated as previously described (4, 7). All mice were backcrossed onto the C57BL/6J background. Littermate controls were all homozygous for the *Mogat1* floxed allele. Mice were group housed and given free access to food and water and subjected to a 12 h light/dark cycle.

Eight-week-old male and female mice were given either a control low-fat diet (LFD, Research Diets, 10 kcal % fat matched sucrose, D12450J) or a high-fat diet (HFD, Research Diets, 60 kcal % fat, D12492) for the duration indicated. Before sacrifice, mice were fasted for 4 h starting at 0900 and were euthanized via CO_2_ asphyxiation. Blood was collected from venipuncture of the inferior vena cava into EDTA-coated tubes and plasma was removed by centrifugation. Adipose and other tissues were immediately collected, flash-frozen in liquid nitrogen, and stored at −80°C until further use.

### Metabolic phenotyping

Glucose tolerance tests (GTT) were performed in 5 h fasted mice with the fasting beginning at 0900. Glucose was dissolved in saline and mice were given intraperitoneal (IP) injections of glucose solution (1 g/kg lean mass). Insulin tolerance tests (ITT) were performed in mice fasting 4 h starting at 0900. Insulin was given as an IP injection of recombinant Humulin R® (0.75 U/kg lean mass in saline). Blood glucose in all studies was measured from a drop of tail blood using a One Touch Ultra glucometer (Life Scan Inc.) at times indicated. Lean and fat mass were determined in fed mice by ECHO MRI.

### Adipocyte sizing and counting

Adipocyte size and number were measured as previously described (13, 14). Briefly, harvested inguinal and gonadal white adipose tissue (25-50 mg) was fixed in 0.2 mol/L collidine HCl/31 mg/mL osmium tetraoxide solution for 30 days followed by dissociation with 8 mol/L urea and 154 mmol/L NaCl. Liberated adipocytes were measured using a 400 micrometer aperture on a Multisizer 3 (Beckman Coulter).

### Plasma analytes

Plasma lipids were determined enzymatically using commercially available kits: triglycerides (Infinity Triglyceride colorimetric assay kit (Thermo Fisher, TR22421), non-esterified free fatty acids (WAKO), and free glycerol reagent (Sigma, F6428).

### Isolation of stromal vascular fractions

Male or female mice were euthanized via CO_2_ asphyxiation and inguinal subcutaneous fat pads were isolated under sterile conditions (15). The resulting fat pads were rinsed in PBS and minced in ice-cold digestion buffer (7.5 mg/ml Collagenase D (Roche 1108886001), and 4.8 mg/ml Dispase II (Sigma D4693) in PBS with 10 mM calcium). Samples were incubated in digestion buffer for 5 min in a 37°C water bath and transferred to an orbital shaker at 37°C / 5% CO_2_ for 30-40 min. Following incubation, cells were passed through a 70 micron filter and centrifuged at 700 x g for 5 min. The resulting stromal vascular fraction was washed in PBS before red blood cells were lysed (ACK lysing buffer, Gibco A10492). The remaining cell pellet was washed twice more with PBS before plating (DMEM F-12, 10% FBS, 100 U/mL PenStrep) and allowed to expand before differentiation or immortalization.

### Immortalization and generating of conditional knockout cells

SVF cells were isolated as above and on the same day a confluent 10 cm dish of 293T cells were transfected with 1.5 µg psPAX2 lentiviral packaging vector (Addgene 12260), 1.5 µg pMD2.G envelop plasmid (Addgene, 12259), and 3 µg pBABE-neo largeT antigen cDNA (Addgene, 1780 ref (16))using lipofectamine 2000 (Invitrogen, 11668019). Three days after the SVF media was replaced with media from the transfected 293T cells that was filtered and mixed at a 1:1 ratio with standard SVF media and viral media with added polybrene (8 µg/mL, Millipore Sigma TR-10030). The infected cells were selected for using Geneticin (G418 Sulfate, 0.2 mg/mL). Conditional knockout cells were generated by infection with media from a confluent 10 cm dish of 293T cells transfected with 3 µg pCL-Eco (Addgene, 12371 ref (17)) and 3 µg MSCV CreERT2 puro (Addgene, 2276 ref (18)) and polybrene (8 µg/mL, Millipore Sigma TR-10030). Cells were selected by puromycin (2 µg/mL) and maintained in DMEM F-12 (10% FBS).

### Differentiation of adipocytes

All cells were allowed to become confluent in 12 well plates. The following day (day 0) cells were treated with a differentiation cocktail mix (DIXγ: 1 µM dexamethasone, 2 µg/mL insulin, 500 µM 3-isobutyl-1-1methylxathine (IBMX), and 15 µM troglitazone). Every 2 days thereafter, the media was replaced with maintenance media (DMEM with 2 µg/mL insulin) for days indicated (5, 6, or 10 days). For knockdown studies, Cre-ERT2 recombinase activity was activated via the addition of 4 nM tamoxifen (TAM) on Day 0 or as otherwise indicated (15). All experiments were performed in triplicate and replicated in at least three independent experiments.

### Oil Red O staining and quantification

Oil Red O staining was performed on days indicated in 12 well plates. Cells were rinsed in PBS and fixed with 10% formalin buffer for 60 min at room temperature. Fixed cells were washed in deionized water and dehydrated with 60% isopropanol for 5 min. Oil Red O was prepared and filtered according to the manufacturer’s suggestions (Sigma O0625). Neutral lipids were stained for 5 min at room temperature, and excess Oil Red O was removed and rinsed in deionized water until clear. Oil Red O was extracted by the addition of 100% isopropanol for 5 min and absorbance was read at 492 nm.

### mRNA isolation and quantitative PCR

Total RNA was isolated from cells or frozen adipose tissue samples using Trizol^TM^ Plus RNA Purification System (Thermo Fischer, 12183555) according to the manufacturer’s protocol. 2 ug RNA was reverse transcribed into cDNA using Taqman high capacity reverse transcriptase (Life Technologies, 43038228). Quantitative PCR was performed using Power SYBR green (Applied Biosystems, 4367659) and measured on an ABI PRISM 7500 or ABI QuantStudio 3 sequence detection system (Applied Biosystems). Results were quantified using the 2^-ΔΔCt^ method and shown as arbitrary units relative to control groups. Primer sequences are listed in Table S1.

### Bulk RNA Sequencing and Analysis

Bulk RNA sequencing and analysis were performed at the Genome Technology Access Center at the McDonnell Genome Institute of Washington University School of Medicine. Samples were prepared according to library kit manufacturer’s protocol, indexed, pooled, and sequenced on an Illumina HiSeq. Basecalls and demultiplexing were performed with Illumina’s bcl2fastq software and a custom python demultiplexing program with a maximum of one mismatch in the indexing read. RNA-seq reads were then aligned to the Ensembl release 76 primary assembly with STAR version 2.5.1a (19). Gene counts were derived from the number of uniquely aligned unambiguous reads by Subread:featureCount version 1.4.6-p5 (20). Isoform expression of known Ensembl transcripts were estimated with Salmon version 0.8.2 (21). Sequencing performance was assessed for the total number of aligned reads, total number of uniquely aligned reads, and features detected. The ribosomal fraction, known junction saturation, and read distribution over known gene models were quantified with RSeQC version 2.6.2 (22).

All gene counts were then imported into the R/Bioconductor package EdgeR (23) and TMM normalization size factors were calculated to adjust for samples for differences in library size. Ribosomal genes and genes not expressed in the smallest group size minus one samples greater than one count per million were excluded from further analysis. The TMM size factors and the matrix of counts were then imported into the R/Bioconductor package Limma (24). Weighted likelihoods based on the observed mean-variance relationship of every gene and sample were then calculated for all samples with the voomWithQualityWeights (25). The performance of all genes was assessed with plots of the residual standard deviation of every gene to their average log-count with a robustly fitted trend line of the residuals. Differential expression analysis was then performed to analyze for differences between conditions and the results were filtered for only those genes with Benjamini-Hochberg false-discovery rate adjusted p-values less than or equal to 0.05.

The heatmap was generated using iDEP 9.0 (26). For each contrast extracted with Limma, global perturbations in known Gene Ontology (GO) terms, MSigDb, and KEGG pathways were detected using the R/Bioconductor package GAGE (27) to test for changes in expression of the reported log 2 fold-changes reported by Limma in each term versus the background log 2 fold-changes of all genes found outside the respective term. Perturbed KEGG pathways where the observed log 2 fold-changes of genes within the term were significantly perturbed in a single-direction versus background or in any direction compared to other genes within a given term with p-values less than or equal to 0.05 were rendered as annotated KEGG graphs with the R/Bioconductor package Pathview (28).

### Lipidomic analysis

Lipidomic analysis was performed at the Washington University Metabolomics Facility. Immortalized SVF cells (5 x 10^6^) were grown in 10 cm dishes and treated with vehicle or DIXγ for 10 days. Following differentiation, cells were pelleted in 500 uL PBS, frozen in liquid nitrogen, and stored at −80°C until analysis. Lipids were extracted in the presence of their internal standards. MAG, DAG, and TAG species were extracted using the Blyth and Dryer method and PA was extracted using protein precipitation before LC-MS/MS analysis. Peak area ratios of each analyte were determined from the internal standards. Quality control (QC) samples were prepared by pooling an aliquot of each study sample and ejected every four study samples. Only species with CV %, 15% for the QC injections were reported. Data are reported as fold change from undifferentiated vehicle-treated cells.

### Statistical analysis

Data were analyzed using GraphPad Prism software. Independent and paired T-tests, one-way analysis of variance (ANOVA), or factorial ANOVAs were performed where appropriate. Secondary post-hoc analysis found differences in groups using either Tukey or Sidak’s multiple comparisons were appropriate. *p* < 0.05 was considered significant.

## RESULTS

### *Mogat1* is required for adipocyte differentiation *in vitro*

We have previously shown *Mogat1* expression and activity are highly induced during differentiation of mouse and human adipocytes *in vitro* (4), but whether this is critical to the differentiation process is unclear. To this end, we isolated stromal vascular fractions (SVF) from inguinal white subcutaneous adipose tissue (iWAT) and differentiated them using a standard differentiation cocktail (DIXγ: 1 µM dexamethasone, 2 µg/mL insulin, 500 µM IBMX, 15 µM troglitazone) (Figure S1A and refs 4, 15). SVFs from wild-type mice had typical increases in adipogenic gene expression that reached maximal levels of expression around day six-post DIXγ treatment (Figure S1B). *Mogat1* expression was induced nearly 100-fold compared to undifferentiated cells, while *Mogat2* expression actually decreased during differentiation (Figure S1C and ref (29)). We found that SVF cells from constitutive *Mogat1* null mice differentiated normally (Figure S2), which is consistent with our previous studies indicating that germline deletion of *Mogat1* does not alter adiposity in mice (7).

To avoid potential compensation of other enzymes with MGAT activity, which are induced during differentiation (Figure S1C), we utilized SVF cells isolated from *Mogat1* floxed mice to generate stable cell lines with an inducible Cre-ERT2 recombinase system (15). Following isolation, the cells were immortalized using SV40 T-antigen and then transfected with an inducible Cre-ERT2 (Figure 1A). After reaching 100% confluency (day-1), the following day (day 0), cells were treated with TAM (or vehicle) and medium containing DIXγ (Figure 1A). Vehicle-treated cells differentiated as expected as indicated by increased Oil Red O staining of neutral lipids (Figure 1B). However, *Mogat1* knockout cells had decreased Oil Red O accumulation (Figure 1B). Importantly, TAM exposure alone is not sufficient to suppress differentiation in *Mogat1* floxed or wild-type SVF cells (Figure S3). As expected, *Mogat1* was highly induced by differentiation in control cells, but TAM treatment reduced differentiation-induced *Mogat1* gene expression without affecting *Mogat2* (Figure 1C). Given the diminished capacity to accumulate lipids with *Mogat1* knockout, we also analyzed the expression of several key regulators of adipocyte differentiation (Figure 2). Expression of *Adipoq*, *Pparg1*, *Pparg2*, *Srebf1*, and *Fasn* were all highly induced during differentiation in control cells (Figure 2A-E). However, the activation of these genes was markedly blunted in the *Mogat1* knockout cells (Figure 2A-E). These data suggest that *Mogat1* expression is indispensable for initiation of adipocyte differentiation of SVF-derived preadipocytes *in vitro*.

**Figure 1:**
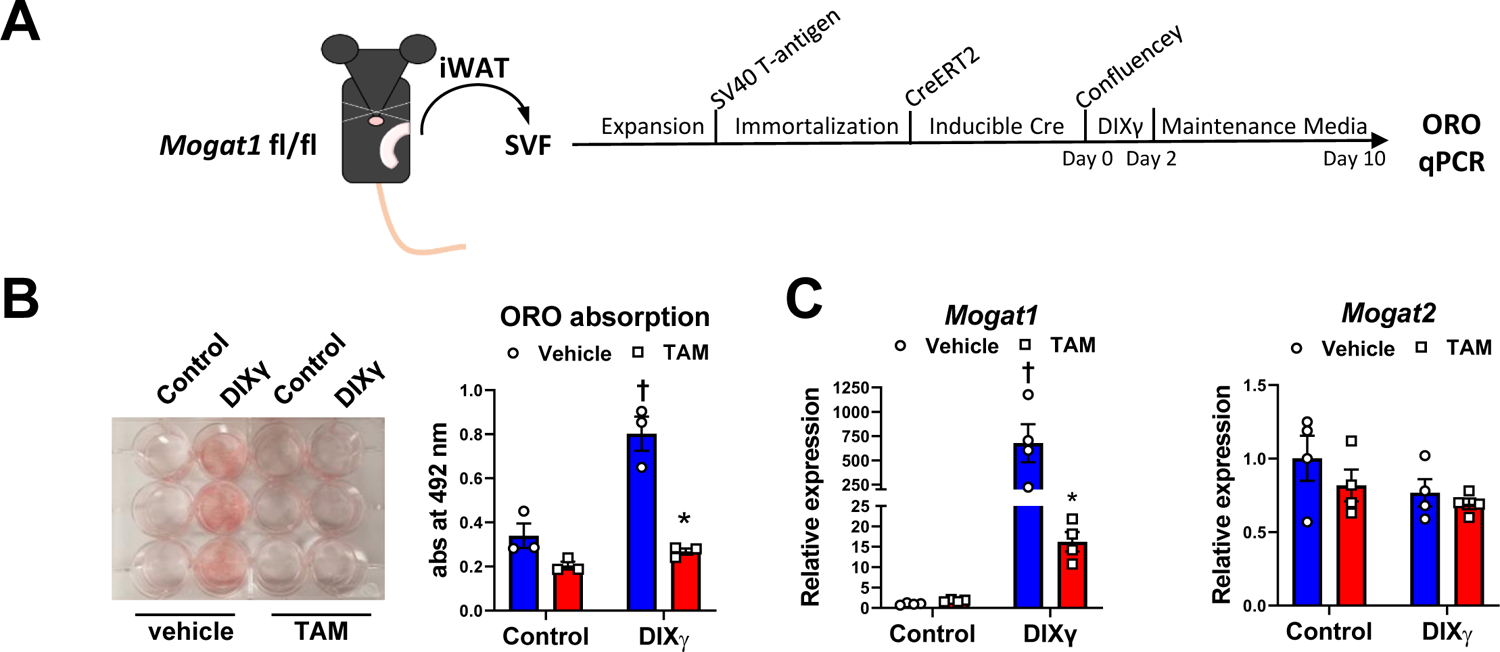
*Mogat1* expression is required for adipocyte differentiation initiation *in vitro.* Stromal vascular fractions (SVF) were isolated from iWAT adipose tissue of *Mogat1* fl/fl mice. Cells were immortalized with SV40 T antigen prior to transfection with an inducible CreERT2. **A:** Diagram of treatment protocol for the generation of SVF and inducible *Mogat1* cell lines as describe in the methods **B, C:** Cre recombinase activity to delete *Mogat1* was activated with the addition 4 nM tamoxifen on Day 0 while other cells were treated with vehicle. **B:** After 10 days of differentiation, lipid accumulation was reduced by *Mogat1* knockout as shown by a representative ORO staining and quantification by extraction and measuring absorption at 492nm. **C:** Gene expression analysis as measured by qPCR. *Mogat1* gene expression is induced by differentiation and inhibited by TAM while *Mogat2* was unaffected by differentiation or TAM treatment. Data are expressed as mean ± S.E.M. * *p* < 0.05, comparisons are indicated; *n* = 3-4 biological replicates.

**Figure 2:**
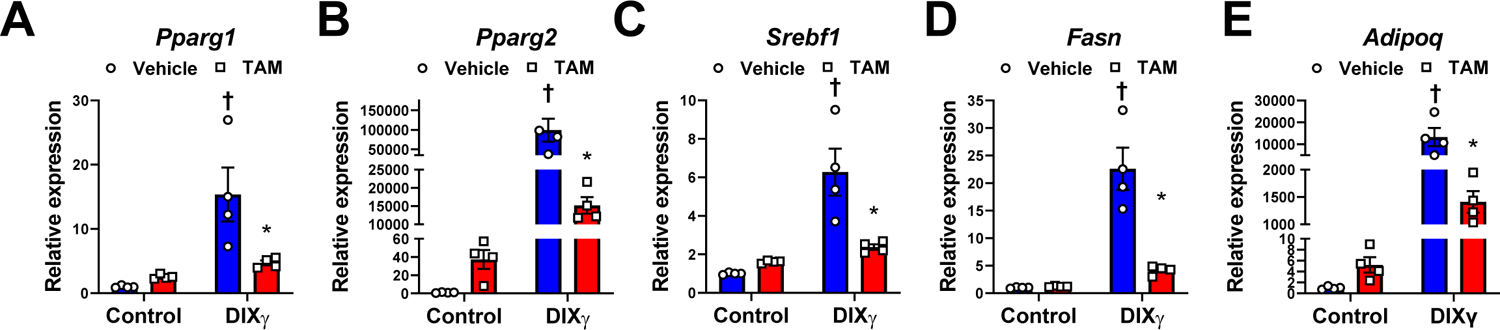
*Mogat1* is required for adipogenic gene expression *in vitro.* Confluent cells were treated with tamoxifen (4 nM) at the onset of differentiation (via DIXγ cocktail) and harvested after 10 days. A-E: Gene expression analysis as measured by qPCR: *Peroxisome proliferator activated receptor gamma (Pparg) 1* and *2, Sterol regulatory element binding transcription factor 1 (Srebf1), Fatty acid synthase (Fasn),* and *Adiponectin (Adipoq)* were increased during differentiation and reduced by *Mogat1* knockout. Data are expressed as mean ± S.E.M. * *p* < 0.05 vs. vehicle, † *p < 0.05 vs.* control undifferentiated cells; *n* = 4 biological replicates.

### *Mogat1* deletion prevents glycerolipid accumulation in differentiated adipocytes

Next, we performed lipidomic analyses to determine the effects of differentiation and *Mogat1* knockout on glycerolipid intermediate abundance (Figure 3A). We found that several MAG species were actually decreased following differentiation and this was unaffected by *Mogat1* knockout (Figure 3B). Interestingly, in wild-type cells that were differentiated, the abundance of MAG(20:4) and MAG(22:6) were extremely low compared to undifferentiated cells (Figure 3B). However, these MAG species were significantly higher in *Mogat1* knockout adipocytes compared to wild-type adipocytes. The product of MGAT activity, DAGs, were increased in differentiated cells compared to undifferentiated cells and many highly abundant DAGs were reduced by *Mogat1* knockout including saturated and monounsaturated DAG(16:0/X) and DAG(16:1/X) species (Figure 3C and Figure S4). Several long-chain highly unsaturated DAGs were decreased during differentiation independent of *Mogat1* expression (Figure S4A, B). TAG species followed a similar trend as DAGs (Figure 3D and Figure S4-C). The DAG precursor and lipin enzymatic substrate, phosphatidic acid (PA), was mostly unchanged by differentiation (Figure 3E), but PA(34:1) and PA(34:2) were decreased in differentiated *Mogat1* knockout cells compared to control cells. Finally, we analyzed the expression of genes that encode glycerolipid metabolic enzymes in these cells. The expression of *monoacylglycerol lipase* (*Mgll*) and both *Dgat1* and *Dgat2* was increased by differentiation and this increase was attenuated by *Mogat1* knockout (Figure 4B, C). Interestingly, in these cells, expression of *Lpin2* but not *Lpin1* increased during differentiation, but was not affected by loss of *Mogat1*, while *Lpin3* expression was increased by *Mogat1* knockout in differentiated cells (Figure 4D). These data suggest that *Mogat1* is required for preadipocytes to fully differentiate and store glycerolipids *in vitro*.

**Figure 3:**
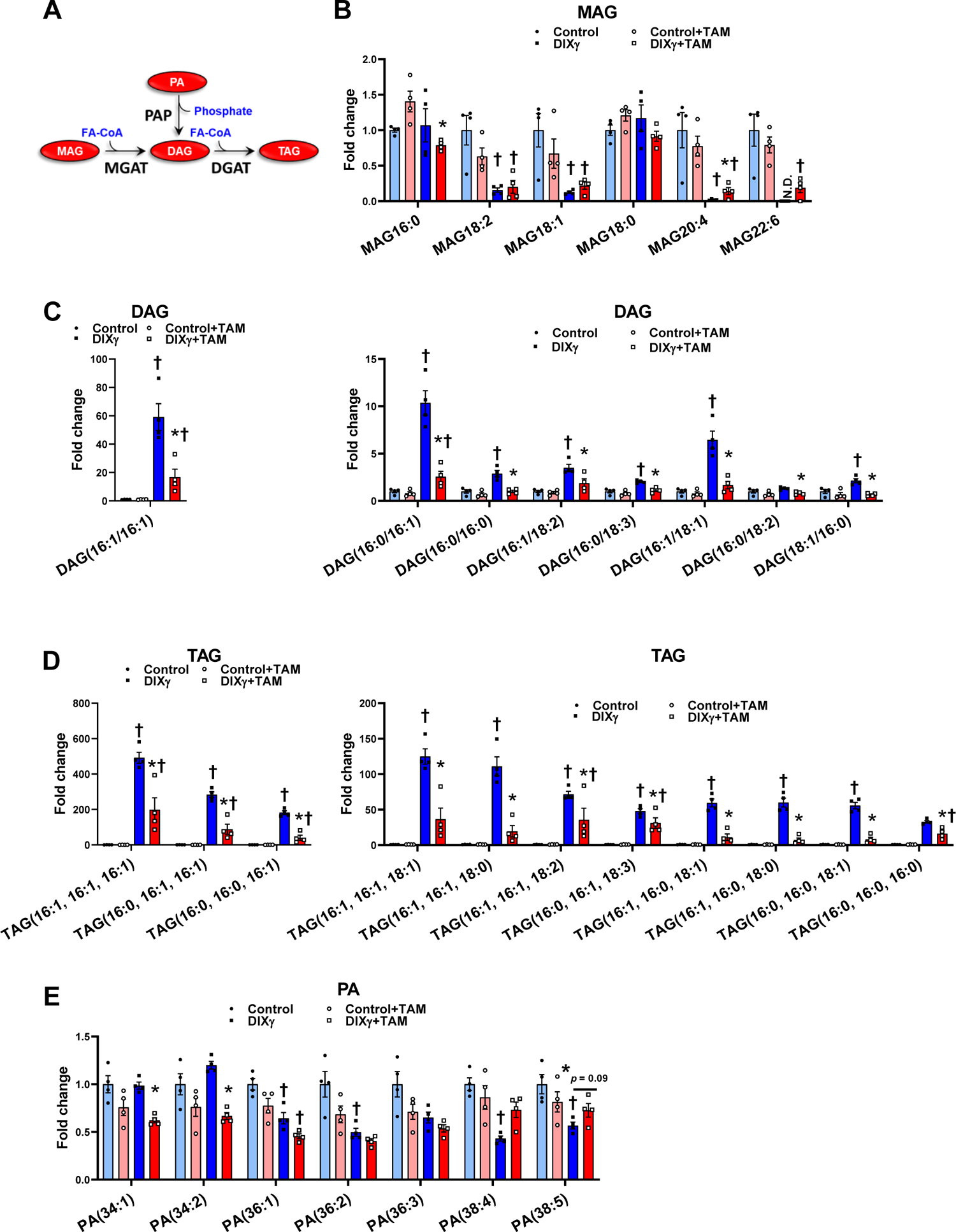
*Mogat1* knockout prevents accumulation of glycerolipids in differentiated SVF cells. Confluent cells were treated with tamoxifen (4 nM) at the onset of differentiation (via DIXγ cocktail) and harvested after 10 days. **A:** Depiction of lipids analyzed via LC-MS/MS **B-E:** Lipid content was lowered by *Mogat1* knockout. Data are expressed as mean ± S.E.M. * *p* < 0.05 vs. vehicle, † *p < 0.05 vs.* control undifferentiated cells; *n* = 4 biological replicates.

**Figure 4:**
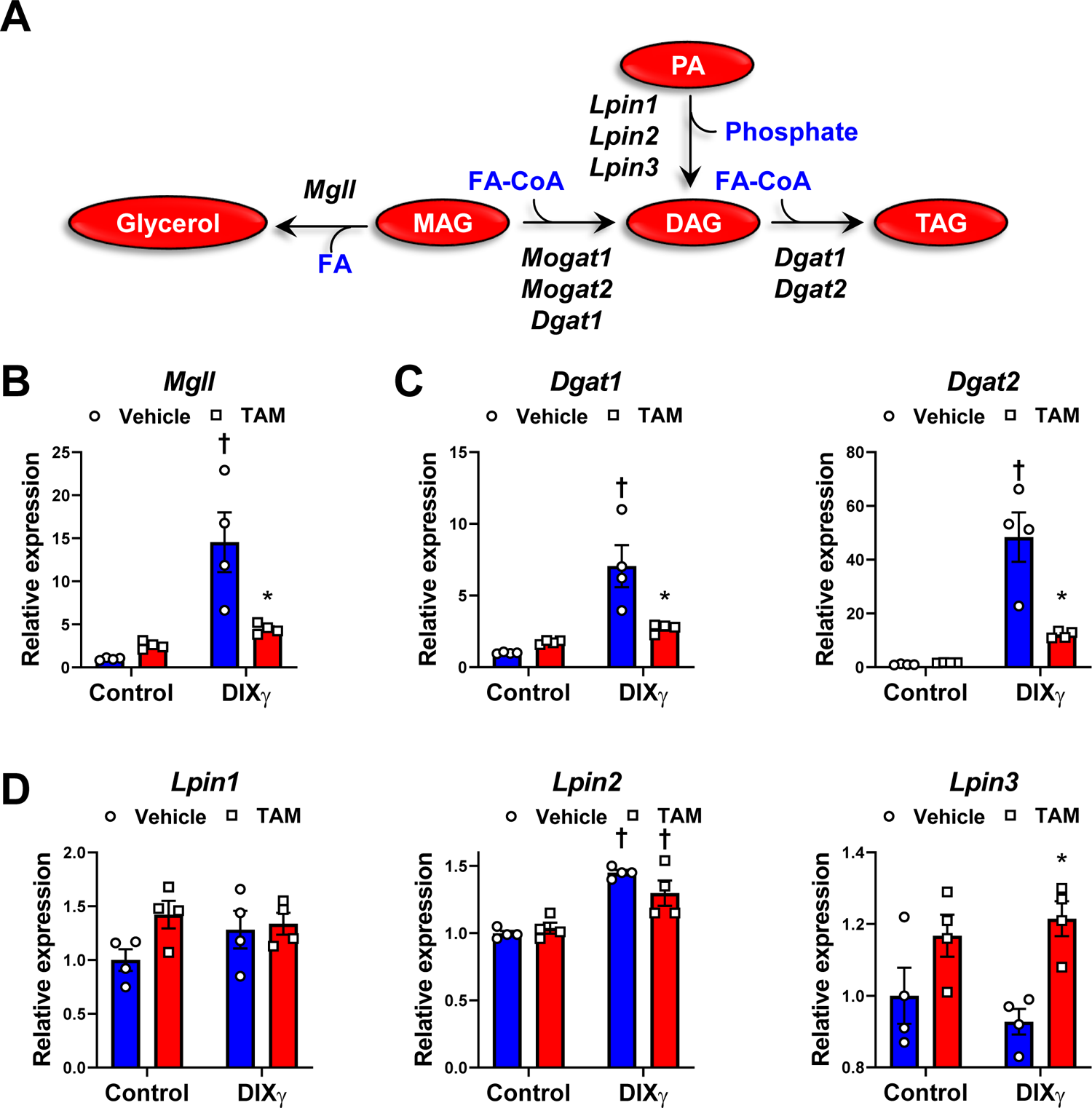
Deletion of *Mogat1* attenuates the induction of triglyceride synthetic enzymes during adipogenesis *in vitro.* Confluent cells were treated with tamoxifen (4 nM) at the onset of differentiation (via DIXγ cocktail) and harvested after 10 days. **A-D:** Gene expression analysis as measured by qPCR. **B:** *Monoacylglycerol lipase (Mgll),* **C:** *Diacylglycerol O-acyltransferase 1 and 2 (Dgat1&2)*. **D:** *Phosphatidate phosphohydrolases (Lpin1,2,&3)*. Data are expressed as mean ± S.E.M. * *p* < 0.05 vs. vehicle, † *p < 0.05 vs.* control undifferentiated cells; *n* = 4 biological replicates.

### Adipocyte-specific *Mogat1* knockout mice gain similar fat mass as littermate controls on a high-fat diet

We have previously reported that, compared to wild-type littermate controls, mice with adipocyte-specific *Mogat1* knockout (Adn-*Mogat1*-/- mice) have reduced fat mass on a chow diet, while whole-body *Mogat1* null mice have increased body weight on HFD (4, 7). To further evaluate the functional role of *Mogat1* in adipose tissue expansion we challenged male and female Adn-*Mogat1*-/- mice and their floxed littermate controls with a HFD (60% kcal from fat) or LFD (10% kcal from fat, matched sucrose) for 16 weeks (Figure 5 and Figure S5). Male Adn-*Mogat1*-/- mice fed HFD gained similar body weight and fat mass on the HFD compared to littermate controls on HFD (Figure 5A, B).

**Figure 5.**
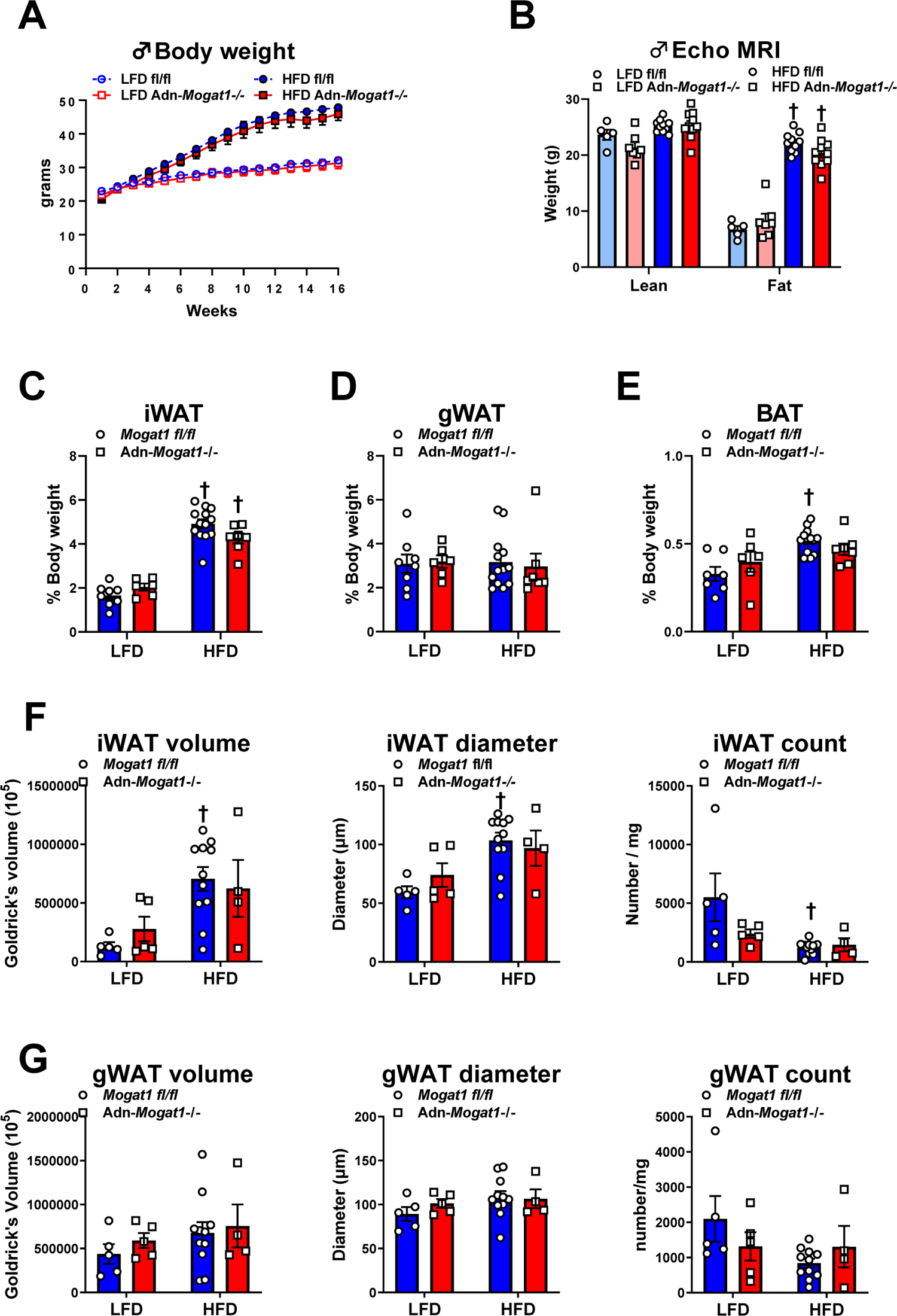
Adipocyte-Specific *Mogat1* knockout mice gain similar weight on a long-term high fat diet compared to littermate controls. Male adipocyte-specific *Mogat1* fl/fl knockout mice (Adn-*Mogat1-/-)* and littermate controls (*Mogat1* fl/fl) were fed a 10% (kcal % fat) low-fat diet (LFD) or a 60% (kcal % fat) high-fat diet (HFD) starting at eight weeks of age. After 16 weeks of diet, mice were fasted for 4 hours prior to sacrifice and tissue collection. **A:** HFD fed mice gained more weight compare to LFD fed controls and was unaffected by *Mogat1* knockout. **B:** After 15 weeks of diet, HFD fed mice had increased fat mass compared to LFD fed mice as measured by Echo MRI. **C-E:** inguinal white adipose tissue (iWAT) and gonadal white adipose tissue (gWAT) but not brown adipose tissue (BAT) were increased by HFD and are expressed as % total body weight. **F, G:** Samples from iWAT and gWAT were fixed and dissociated. Liberated adipocytes were measured and counted. Data are expressed as mean ± S.E.M. \**p* < 0.05 vs. LFD; † *p*, 0.05 vs. *Mogat1* fl/fl mice; *n* = 5-12 in **A-E** and 4-11 in **F,G**.

Individual fat pad mass was also unaffected by genotype (Figure 5C-E). Similar results were found in female mice (Figure S5A-C). WAT samples were fixed and dissociated, and subsequently liberated adipocyte sizes were measured by a particle counter. The volume and diameter of individual iWAT adipocytes were increased by the HFD in wild-type mice but HFD did not affect adipocyte size significantly in the Adn-*Mogat1-/-* mice (Figure 5F). The number of iWAT adipocytes per mg tissue was decreased in HFD wild-types littermate controls (Figure 5F). There were no differences in cell volume, diameter, or number per mg tissue of gWAT between groups (Figure 5G). These data indicate adipocyte *Mogat1* expression does not affect the adiposity or adipocyte size and number in mice fed a HFD.

### Adipocyte-specific *Mogat1* knockout does not improve glucose or insulin tolerance

We have previously shown that *Mogat1* regulates basal lipolysis in adipose explants and 3T3-L1 cells suggesting that loss of *Mogat1* in adipocytes could lead to increased plasma lipids (4). However, plasma TAG, FFA, and glycerol were not different between genotypes on either diet (Figure 6A-C).

**Figure 6.**
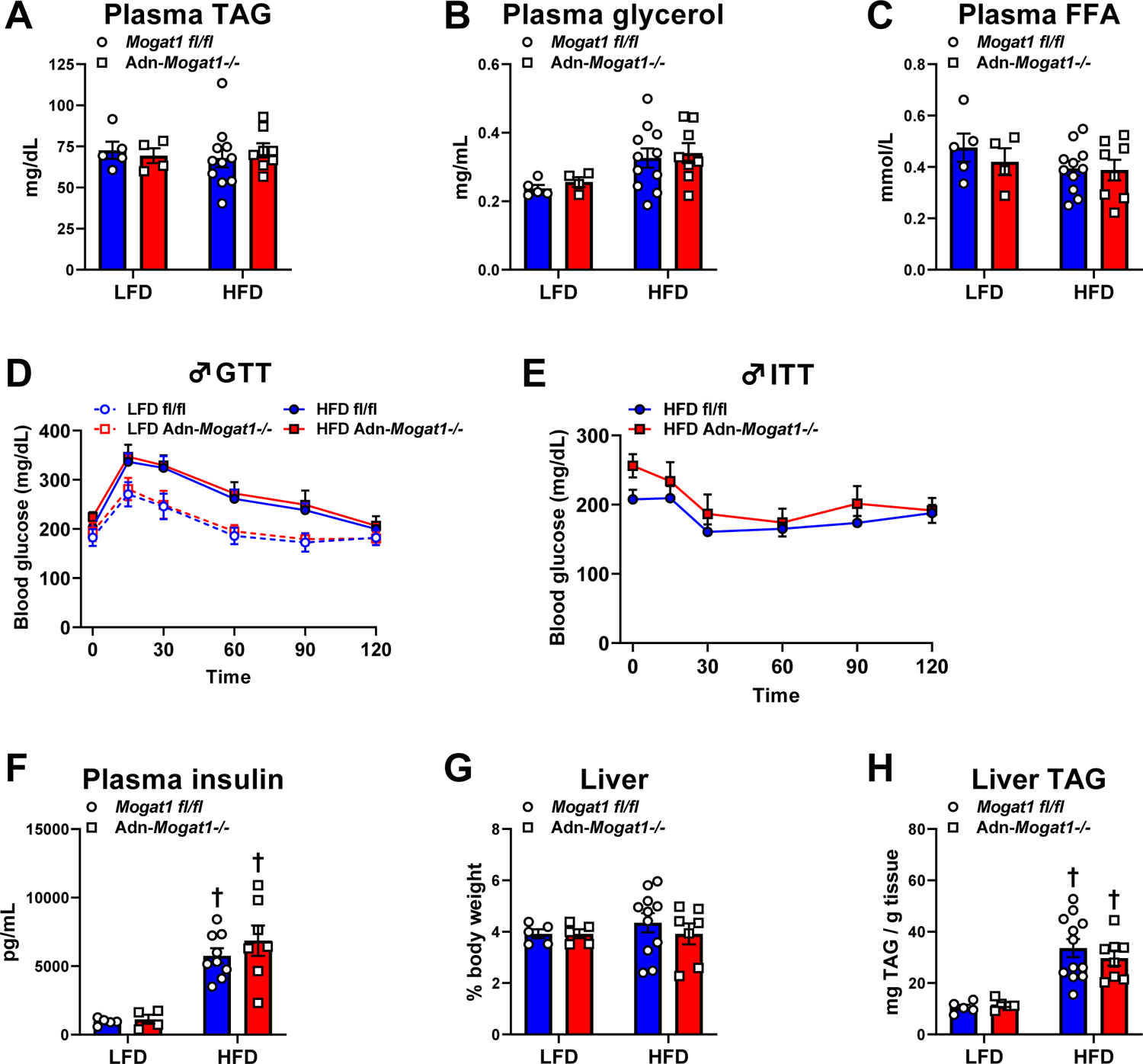
Adn-*Mogat1-/-* mice have a similar metabolic profile as littermate controls. Male Adn-*Mogat1-/-* and littermate control mice were fed a 10% LFD or a 60% HFD starting at eight weeks of age. After 16 weeks of diet, mice were fasted for 4 hours prior to sacrifice and tissue collection. A-C: Plasma triglycerides (TAG), glycerol, and free fatty acids (FFA) were measured enzymatically using commercially available colorimetric assays. D, E: Glucose tolerance tests (GTT, 1 g/Kg lean mass, 5 hour fast) and insulin tolerance tests (ITT, 0.75 U/Kg lean mass, 4 hour fast) were measured at 16 weeks and 15 weeks and were similar between groups. F: Four-hour fasted plasma insulin levels measured at sacrifice were increased by HFD feeding. G: Liver weight as a % total body weight was not changed by diet or genotype. H: Liver triglycerides (TAG) were measured as described above and were increased by HFD feeding. Data are expressed as mean ± S.E.M. †*p* < 0.05 vs. LFD; *n* = 4-12.

We performed glucose and insulin tolerance tests in adipocyte-specific *Mogat1* knockout mice and littermate controls. HFD feeding lead to glucose and insulin intolerance, but there was no difference in tolerance between genotypes on either diet (Figure 6D, E). Likewise, plasma insulin concentrations were increased by HFD, but similar between groups (Figure 6F). Analogous results were observed in female mice fed a HFD (Figure S5D, E). Liver weight did not change but HFD increased hepatic TAG content in both genotypes (Figure 6G, H). These data indicate adipocyte-specific *Mogat1* knockout mice have similar adiposity in response to a HFD as the littermate controls and adipocyte *Mogat1* expression does not affect systemic lipid or glucose metabolism.

### Transcriptomic analysis of inguinal white adipose tissue from adipocyte-specific *Mogat1* knockout mice reveals few differences between genotypes

We first assessed the expression of several lipogenic enzymes using RNA from iWAT from mice on either low fat or HFD. We did not detect any significant increase in the expression of DAG or TAG synthesizing genes (*Mogat2, Dgat1, Dgat2,* and *Lpin1*) (Figure 7A) that might compensate for the lack of *Mogat1* between genotypes (Figure 7B and refs (1, 6, 7, 30, 31). However, *Mogat2* was increased nearly forty-fold by high-fat diet in both genotypes. Expression of classic markers of adipocyte lipogenesis were also unaffected by *Mogat1* deletion except for fatty acid synthase (*Fasn*) which was increased in LFD-fed *Mogat1* knockout mice compared to littermate controls. (Figure 7C).

**Figure 7.**
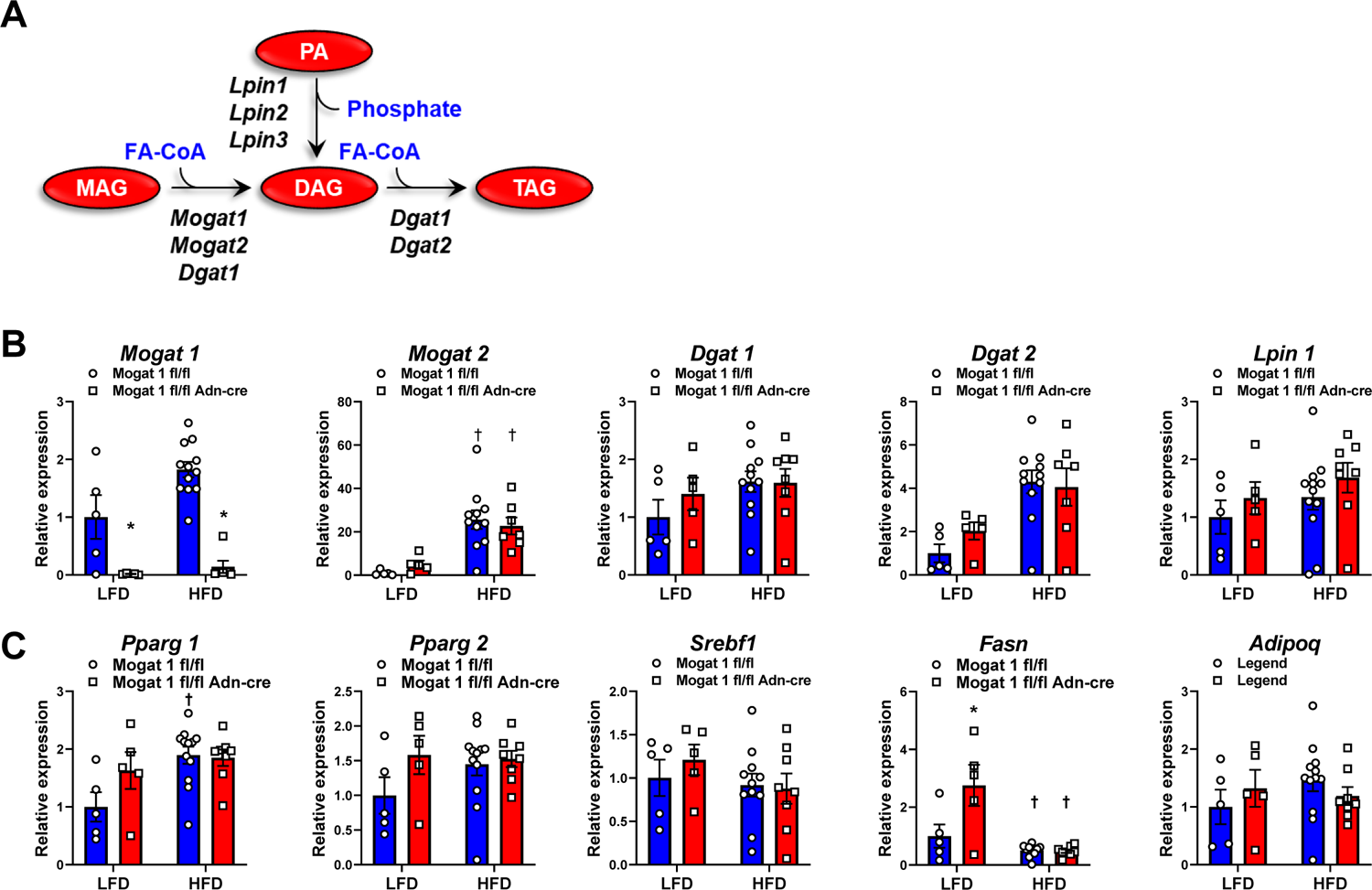
Subcutaneous adipose tissue from adipocyte-specific *Mogat1* knockout mice have a similar gene expression profile as littermate controls. Male Mogat1 fl/fl and littermate adiponectin Cre+ (Adn-*Mogat1*-/-) mice were fed a 10% low fat diet (LFD) or a 60% (HFD) starting at eight weeks of age. After 16 weeks of diet, mice were fasted for 4 hours prior to sacrifice and tissue collection. A schematic of genes involved in TAG metabolism. B, C: Subcutaneous iWAT adipose tissue gene expression analysis as measured by qPCR. A: *Mogat1, Mogat2, Dgat1* and *2,* and *Lpin1.* B: *Peroxisome proliferator activated receptor gamma (Pparg) 1* and *2, Sterol regulatory element binding transcription factor 1 (Srebf1), Fatty acid synthase (Fasn),* and *Adiponectin (Adipoq)* Data are expressed as mean ± S.E.M. * *p* < 0.05 gene effect, † *p* < 0.05 diet effect; *n* = 5-11.

To better understand the molecular pathways affected by adipocyte-specific *Mogat1* deletion, we performed bulk RNA sequencing on iWAT from Adn-*Mogat1*-/- mice and their littermate controls after 16 weeks on HFD. This analysis revealed very few differences between genotypes (Figure S6). In fact, only *Mogat1* was significantly decreased and 5 genes significantly increased given our criteria (adjusted *p-*value < 0.05, log_2_ FC < 2.0, Figure S6B, C). Pathway analysis identified several significantly downregulated GO biological process pathways including those related to fatty acid metabolism and oxidation (Figure S7A, B). While KEGG pathways suggest alterations in metabolic pathways, insulin signaling, and oxidative phosphorylation among others (Figure S7C, D). Although these pathways do not translate to any physiological differences seen in these mice (Figures 5, S5, and 6).

## DISCUSSION

Esterification and storage of free fatty acids as neutral TAG in adipocytes is vital for preserving systemic metabolic health. Multiple enzymatic pathways have evolved to synthesize TAG including the MGAT pathway (Figure 4A). MGAT activity and function have been extensively characterized in intestine (2, 8, 30, 32) and liver (5, 7, 11, 12, 31, 33–35). However, few studies have evaluated MGAT function in adipocytes. Here we demonstrate that *Mogat1* is dispensable for adipose tissue expansion in high-fat-fed mice. We also confirmed our previous report that *Mogat1* is the predominant MGAT isoform enriched during *in vitro* differentiation of mouse preadipocytes (Figure S1 and ref. (4)). Using preadipocytes harboring the conditional *Mogat1* allele, we generated knockout adipocyte cell lines and found that *Mogat1* expression is vital for their differentiation into adipocytes. *Mogat1* knockout reduced the accumulation of neutral lipids and the expression of pro-adipogenic genes like *Pparg1, Pparg2, Srebf1,* and *Fasn.* Lastly, *Mogat1* ablation prevented the accumulation of lipids including a including the most abundant DAG and TAG species during differentiated mouse adipocytes.

We previously demonstrated that *Mogat1* is dispensable for hepatic MGAT activity and steatosis development in obese mice (7). These data suggested compensation by alternative enzymes with MGAT activity and/or alternative sources for cellular TAG. In the present study, the loss of *Mogat1* in adipocytes does not alter adiposity (i.e. TAG accumulation) in diet-induced obese mice. This may not be surprising given the functional redundancy of the lipid synthetic pathways. Lessons from *in vitro* studies imply that the acyltransferases, DGAT1 and DGAT2, are functionally redundant and compensate for the loss of either isoform in genetic deletion studies. The knockout of both DGAT isoforms is required to inhibit *in vitro* adipocyte differentiation (36). In adipose tissue specifically, DGAT1 completely compensates in adipose-specific DGAT2 knockout mice, while DGAT1 knockout mice are modestly leaner than their wild-type counterparts (37, 38). Furthermore, in the present study we found that the adipose tissue expression of *Mogat2*, which is normally very low, is markedly increased by HFD feeding in mice (Figure 7B). This is important given that MGAT2 has a higher specific activity than MGAT1 (1). It is possible that the activity of MGAT2 and other enzymes capable of mediating TAG synthesis overcomes any effect of *Mogat1* deficiency.

Constitutive *Mogat1* deletion does not alter adipocyte differentiation both *in vivo* and *in vitro*, yet acute loss at the onset of differentiation *in vitro* almost completely prevents glycerolipid accumulation and activation of the differentiation program. In addition, the adiponectin-Cre-driven deletion, which occurs after differentiation has commenced, does not affect differentiation *in vivo*. This could suggest *Mogat1* plays a critical role at a very specific early time in the differentiation program and that chronic loss of *Mogat1* results in compensatory effects. Although we have not identified a specific enzyme with compensatory activity, lipin family proteins also generate DAG. Although MGATs and lipins converge on DAG synthesis (the penultimate step in TAG synthesis) the physiological consequences of inhibiting these pathways are dissimilar *in vitro* and *in vivo* (2–4, 39–41). In contrast to prior work in other adipogenic cells (29, 42–44), *Lpin1* and *Lpin3* expression is only modestly increased during our differentiation of the SVF-derived cell lines (Figure 4D). Genetic loss of *Lpin1* leads to dramatic impairments in adipogenesis both *in vitro* and *in vivo* (45, 41, 43, 42). This suggests the possibility that the failure of this particular *in vitro* model to increase *Lpin1* expression during differentiation might explain the dramatic reliance on *Mogat1* in this context and the differences with the *in vivo* phenotype.

Our lipidomics analysis revealed that the most abundant DAG and TAG species accumulate with differentiation and are reduced by *Mogat1* knockout (Figure 3 and Figure S3). In contrast, many MAG and PA species were reduced during differentiation suggesting that these precursors are consumed to drive DAG and TAG synthesis. Lipidomic analysis has increased the breadth of knowledge regarding the importance of specific fatty acyl chains of glycero/phospholipids in metabolic regulation. Specific to this study, both arachidonic acid (20:4) and docosahexaenoic acid (22:6) have been shown to inhibit preadipocyte differentiation *in vitro* (46–48). In the present studies, MAG(20:4) and MAG(22:6) were abundant in undifferentiated cells, but not in differentiated wild-type adipocytes. The reduction that occurred during differentiation was attenuated by *Mogat1* ablation. Yen and colleagues have reported that MGAT1 activity is higher for long-chain highly unsaturated MAGs, specifically MAG(20:4) (1).

Whether these MAG species can directly inhibit adipocyte differentiation remains to be explored. We acknowledge these studies are limited as our lipidomics data represent static lipid abundance and not flux into glycerolipid synthesis. Thus, these differences may be indicative of impaired differentiation and not MGAT1 enzymatic function *per se*.

## CONCLUSION

Here we have provided evidence that *Mogat1* expression initiates adipocyte differentiation in mouse iWAT preadipocytes. Furthermore, *Mogat1* knockout reduces adipogenic gene expression and glycerolipid accumulation in differentiated cells. HFD-fed adipocyte-specific *Mogat1* knockout mice have no discernable metabolic phenotype compared to littermate controls. These data suggest that there are compensatory mechanisms for loss of *Mogat1* and/or that adipocyte *Mogat1* expression is dispensable for diet-induced obesity in mice.

## ACKNOWLEDGMENTS

We thank the Genome Technology Access Center in the Department of Genetics at Washington University School of Medicine for help with genomic analysis. The Center is partially supported by NCI Cancer Center Support Grant #P30 CA91842 to the Siteman Cancer Center and by ICTS/CTSA Grant# UL1 TR000448 from the National Center for Research Resources (NCRR), a component of the National Institutes of Health (NIH), and NIH Roadmap for Medical Research. Mass spectrometry was performed in the Metabolomics Facility at Washington University (P30 DK020579). This publication is solely the responsibility of the authors and does not necessarily represent the official view of NCRR or NIH. We would like to acknowledge the following investigators for their plasmid contributions to Addgene that were used in this research: Didier Trono, Bob Weinberg, Inder Verma, and Tyler Jacks.

---

**Figure S1: Gene expression profiles of the iWAT SVF adipocyte differentiation protocol.** A: Schematic of isolation and treatment protocol for the generation of SVF cell-derived adipocytes from iWAT as describe in the methods. B, C: qPCR showing classical adipocyte markers ad glycerolipid gene expression during differentiation. Data are expressed as mean ± S.E.M. † *p* < 0.05, vs Day 0; *n* = 5 biological replicates.

**Figure S2: *Mogat1-/-* SVF cells differentiated similar to wild-type controls. A:** Oil Red O staining shows similar neutral lipid accumulation in both genotypes. B: qPCR showing classical adipocyte markers ad glycerolipid gene expression during differentiation. Data are expressed as mean ± S.E.M. † *p* < 0.05, vs Day 0; Oil Red O *n* = 2 biological replicates, qPCR n = 3 biological replicates.

**Figure S3: Tamoxifen treatment does not affect differentiation of wild type cells.** SVF cells were differentiated as described and treated with TAM on day indicated. After 10 days of differentiation, lipid accumulation was measured by ORO staining and quantification by extraction and measuring absorption at 492nm.

**Figure S4: *Mogat1* knockout prevents accumulation of glycerolipids in differentiated SVF cells.** Stromal vascular fractions (SVF) were isolated from iWAT of *Mogat1* fl/fl mice. Confluent cells were treated with tamoxifen (4 nM) at the onset of differentiation (via DIXγ cocktail) and harvested after 10 days. A, B: DAG content was analyzed via LC-MS/MS and was lowered by *Mogat1* knockout. Data are expressed as mean ± S.E.M. * *p* < 0.05 vs. vehicle, † *p = 0.05 vs.* control undifferentiated cells; *n* = 4 biological replicates.

**Figure S5. Female Adipocyte-Specific *Mogat1* knockout mice gain similar weight on a long-term high fat diet (HFD) compared to littermate controls.** Female *Mogat1* fl/fl and littermate adiponectin Cre + (Adn-*Mogat1*-/-) mice were fed a 60% HFD starting at eight weeks of age. After 16 weeks of diet, mice were fasted for 4 hours prior to sacrifice and tissue collection. A, B: Weight gain and % lean or fat mass was unaffected by *Mogat1* knockout as measured by Echo MRI. C: gWAT and iWAT but not BAT were increased by HFD and are expressed as % total body weight. D, E: Insulin tolerance tests (ITT, 0.75 U/Kg lean mass, 4 hour fast) and glucose tolerance tests (GTT, 1 g/Kg lean mass, 5 hour fast) were similar between groups. Data are expressed as mean ± S.E.M. n = 7-9; n = 3-4 for ECHO MRI.

**Figure S6. Bulk RNA sequencing analysis of iWAT shows few differences between *Mogat1-/-* and littermate controls on a HFD.** A: Heatmap of merged differential expression data from iWAT from *Mogat1-/-* littermate controls on a HFD. B: Volcano plots of merged differential expression data was graphed as log_2_ fold change verses –log_10_ unadjusted *p* value. C: List of significantly changed genes. Gene expression changes were considered meaningful if they were > log_2_ fold change and had an adjusted *p* value of <0.05 vs. fl/fl mice on HFD.

**Figure S7: *Mogat1* ablation in adipocytes alters metabolic pathways.** A-D: Graphical representation of pathway analysis showing top 10 significant signal direction changes of GO biological process and KEGG Signaling and Metabolism pathways. The color of the pathway label indicates the *p* value and the x-axis is the mean log fold change.

**Table S1.**
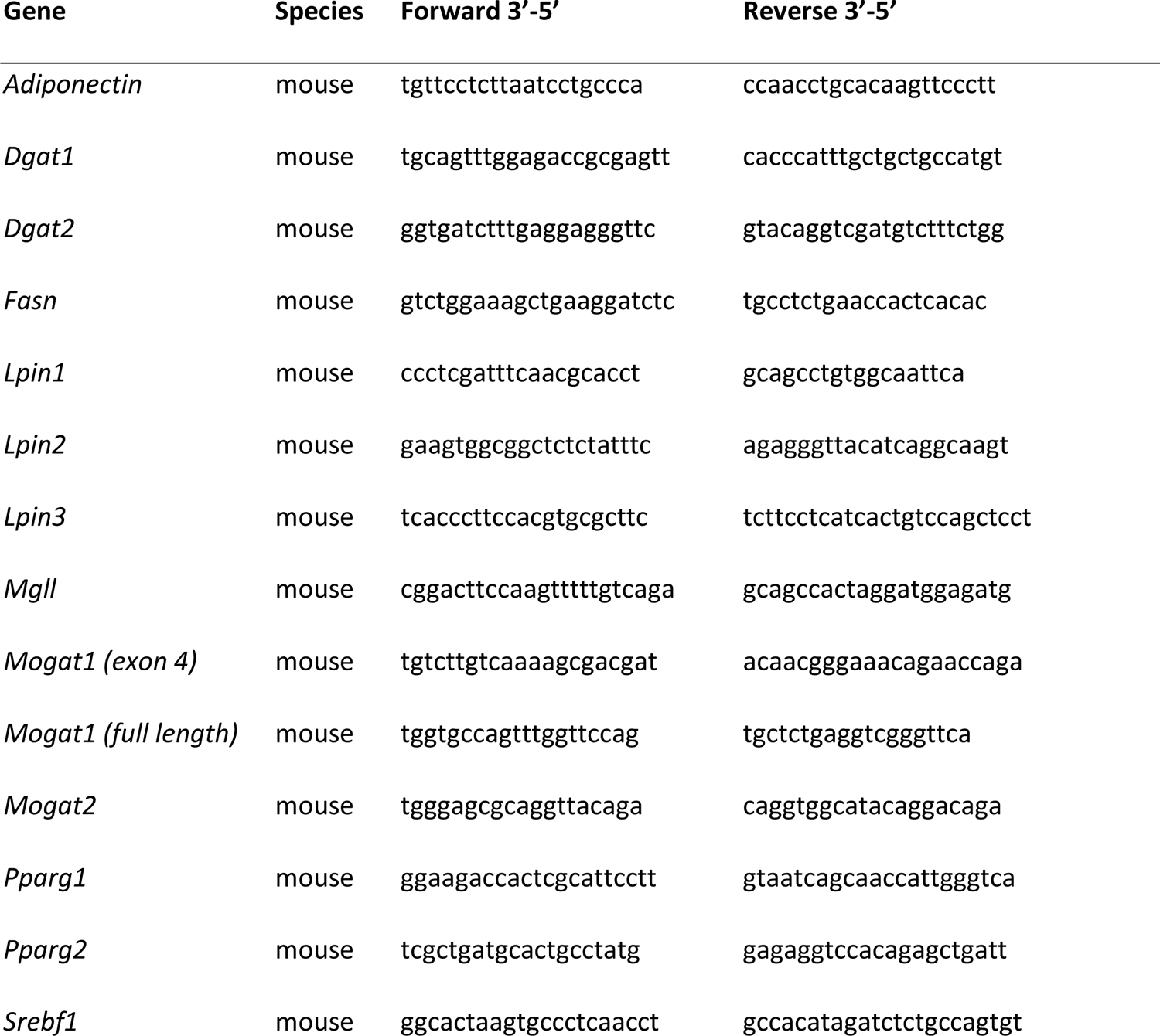
Primer sequences used for RT-qPCR

## Notes

Funding: This work was supported by grants from the NIH R56 DK111735 and the American Diabetes Association 1-17-IBS-109 to BNF, NIH K01 DK126990 to AJL and core laboratories of Washington University School of Medicine: Digestive Diseases Research Cores Center NIH P30DK052574, Nutrition Obesity Research Center P30 DK056341 and the Diabetes Research Center NIH P30 DK020579.

### Competing Interest Statement

The authors have declared no competing interest.

