## Supplemental figures for "Monoacylglycerol O-acyltransferase 1 is required for adipocyte differentiation *in vitro* but does not affect adiposity in mice"

Figure S1

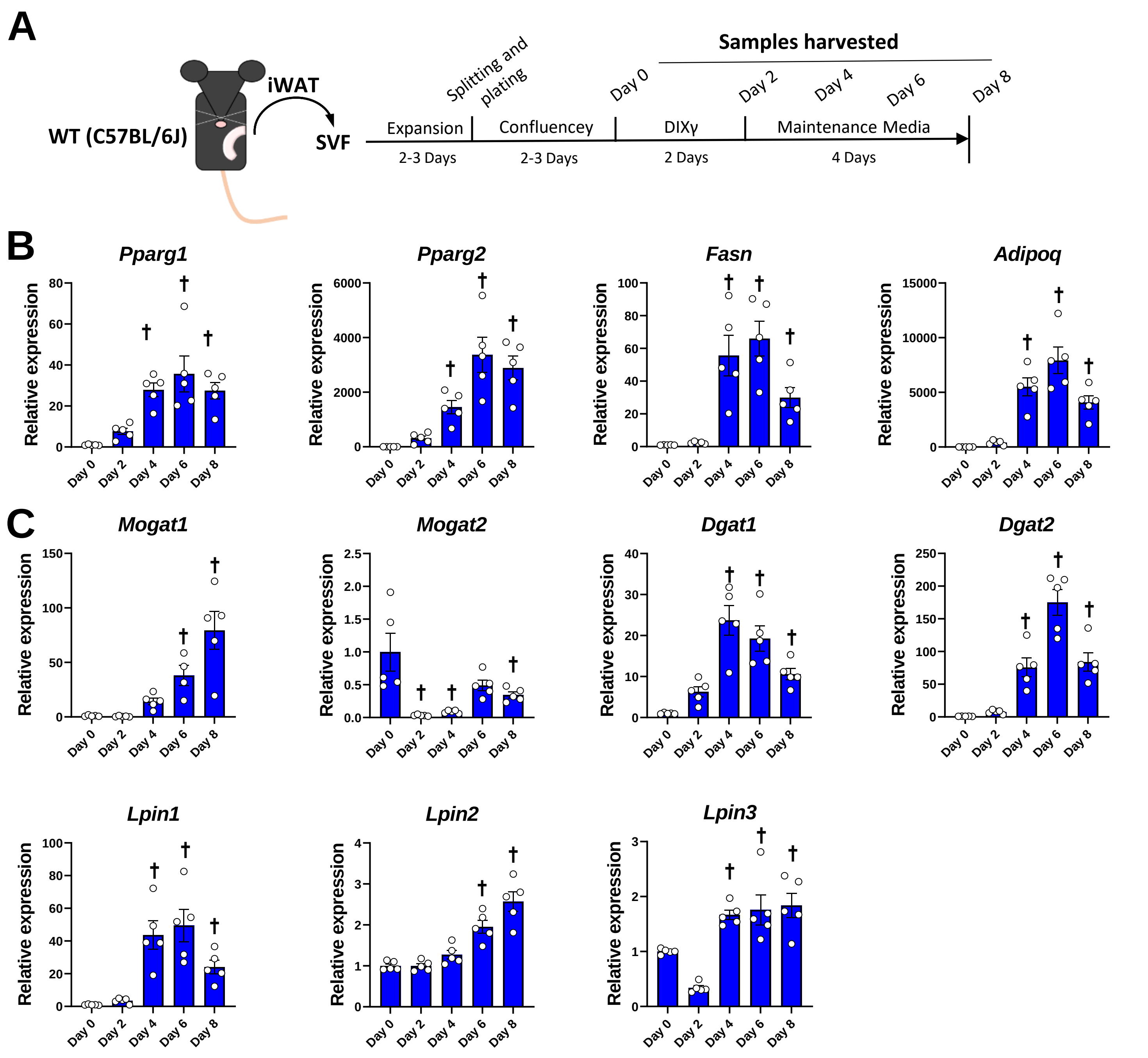


**Figure S1: Gene expression profiles of the iWAT SVF adipocyte differentiation protocol. A:** Schematic of isolation and treatment protocol for the generation of SVF cell-derived adipocytes from iWAT as describe in the methods. **B, C:** qPCR showing classical adipocyte markers ad glycerolipid gene expression during differentiation. Data are expressed as mean ± S.E.M. † $p < 0.05$, vs Day 0; $n = 5$ biological replicates.

### Figure S2

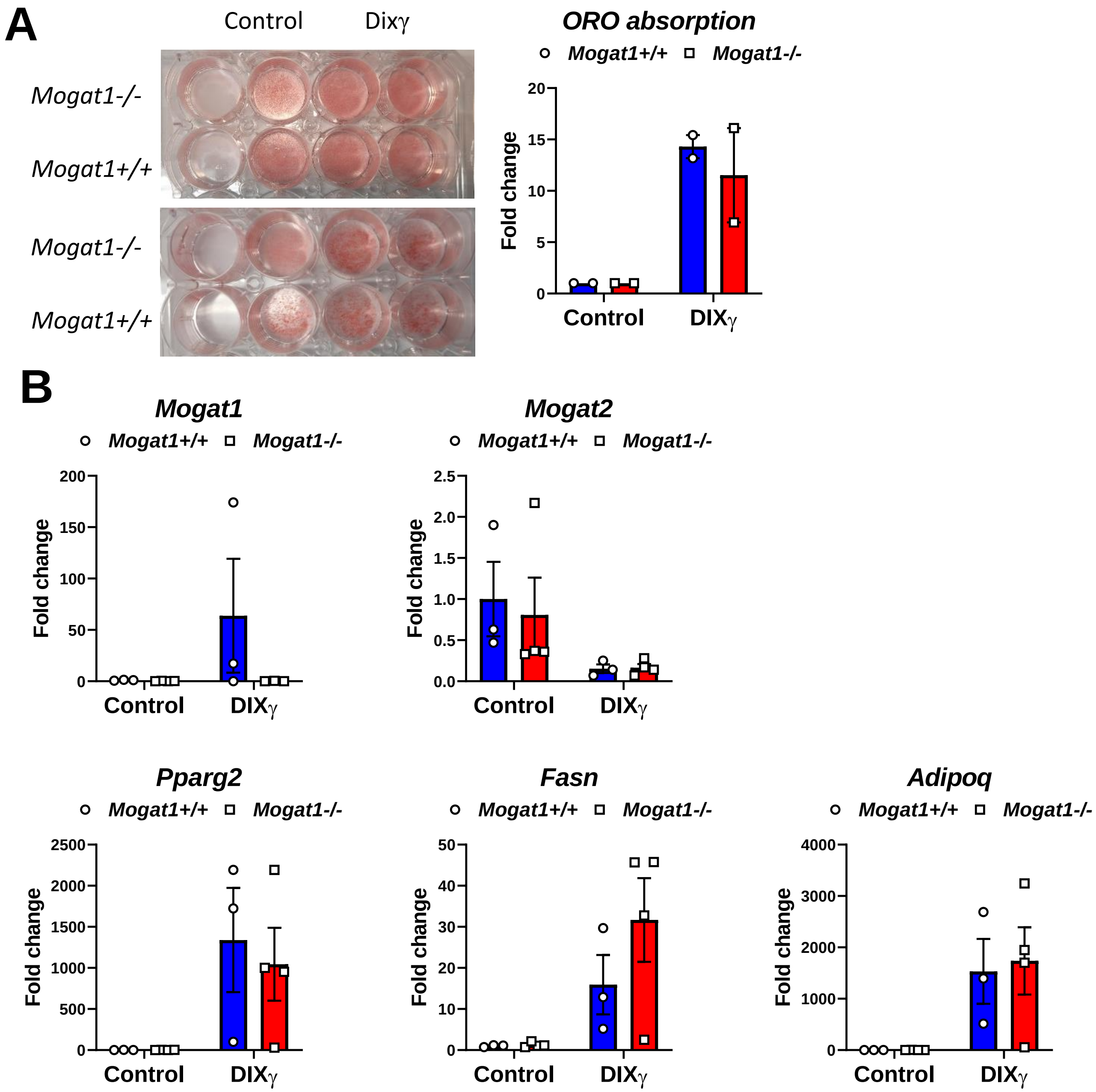


**Figure S2: *Mogat1-/-* SVF cells differentiated similar to wild-type controls. A:** Oil Red O staining shows similar neutral lipid accumulation in both genotypes. **B:** qPCR showing classical adipocyte markers ad glycerolipid gene expression during differentiation. Data are expressed as mean ± S.E.M. † $p < 0.05$, vs Day 0; Oil Red O $n = 2$ biological replicates, qPCR n = 3 biological replicates.

Figure S3

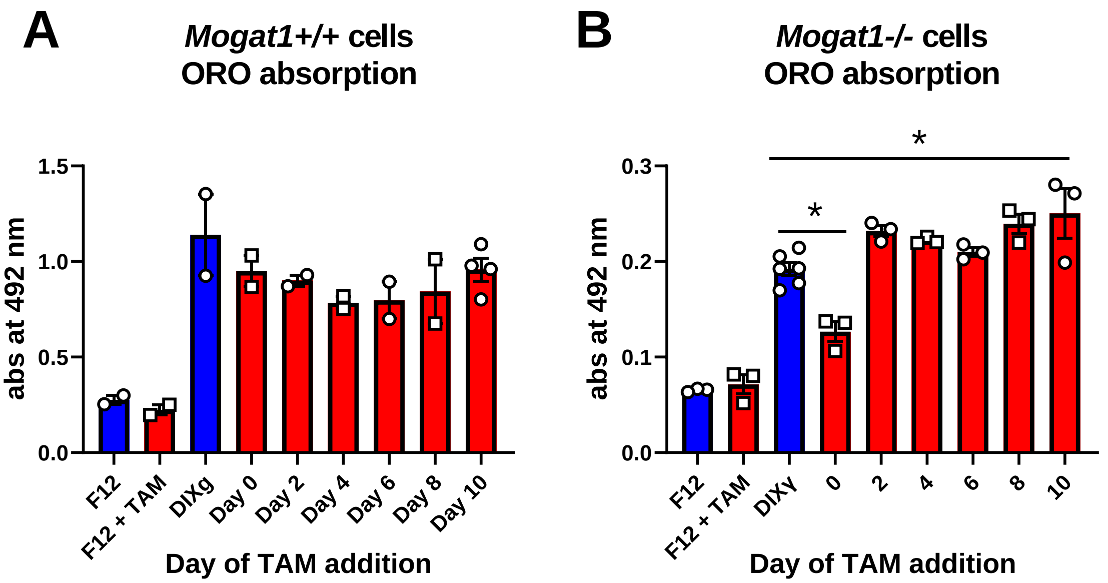


**Figure S3: Tamoxifen treatment does not affect differentiation of wild type cells.** SVF cells were differentiated as described and treated with TAM on day indicated. After 10 days of differentiation, lipid accumulation was measured by ORO staining and quantification by extraction and measuring absorption at 492nm.

### Figure S4

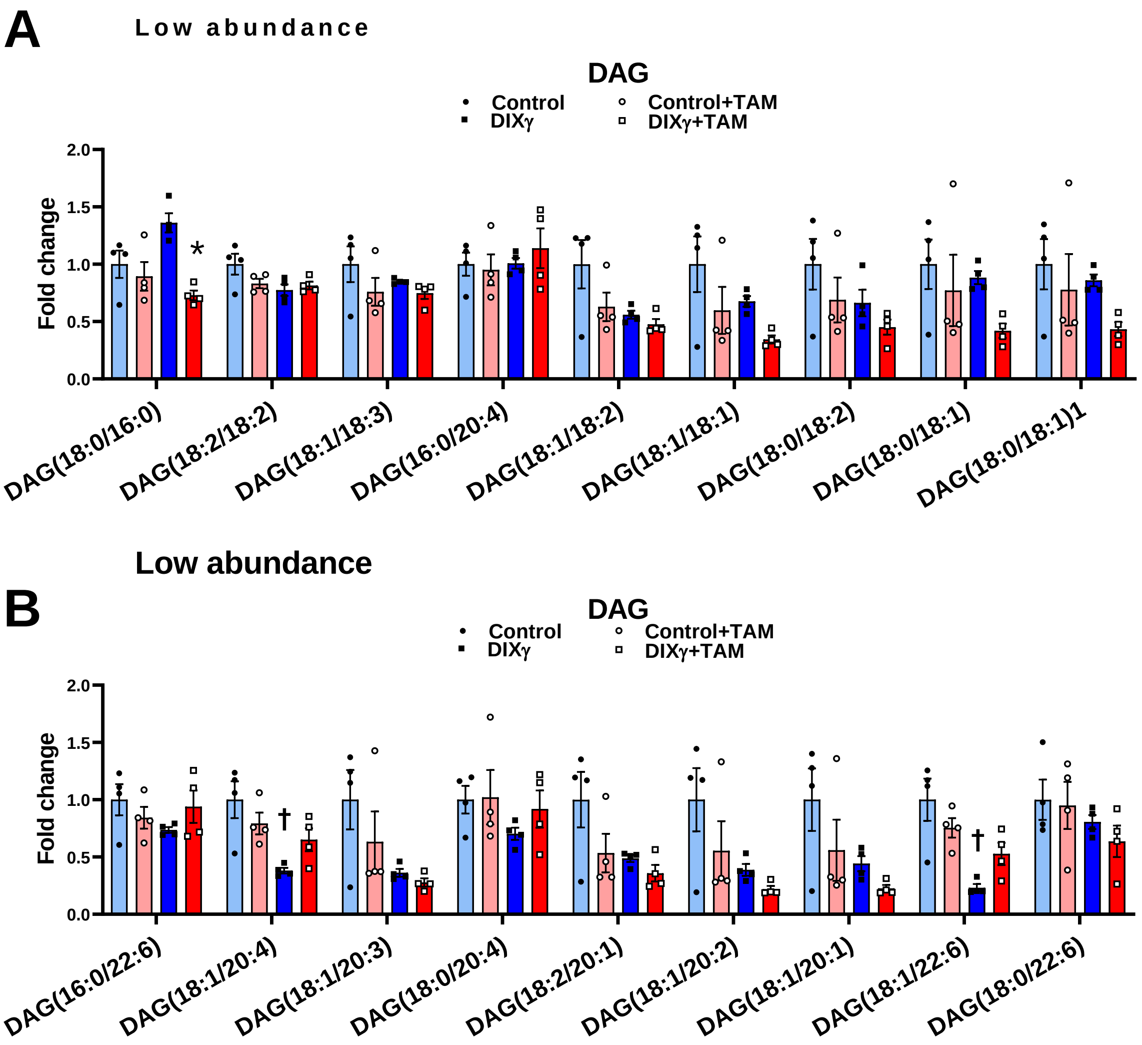


**Figure S4: *Mogat1* knockout prevents accumulation of glycerolipids in differentiated SVF cells.** Stromal vascular fractions (SVF) were isolated from iWAT of *Mogat1* fl/fl mice. Confluent cells were treated with tamoxifen (4 nM) at the onset of differentiation (via DIXγ cocktail) and harvested after 10 days. **A, B:** DAG content was analyzed via LC-MS/MS and was lowered by *Mogat1* knockout. Data are expressed as mean ± S.E.M. * $p < 0.05$ vs. vehicle, † $p < 0.05$ *vs.* control undifferentiated cells; $n = 4$ biological replicates.

### Figure S4 continued

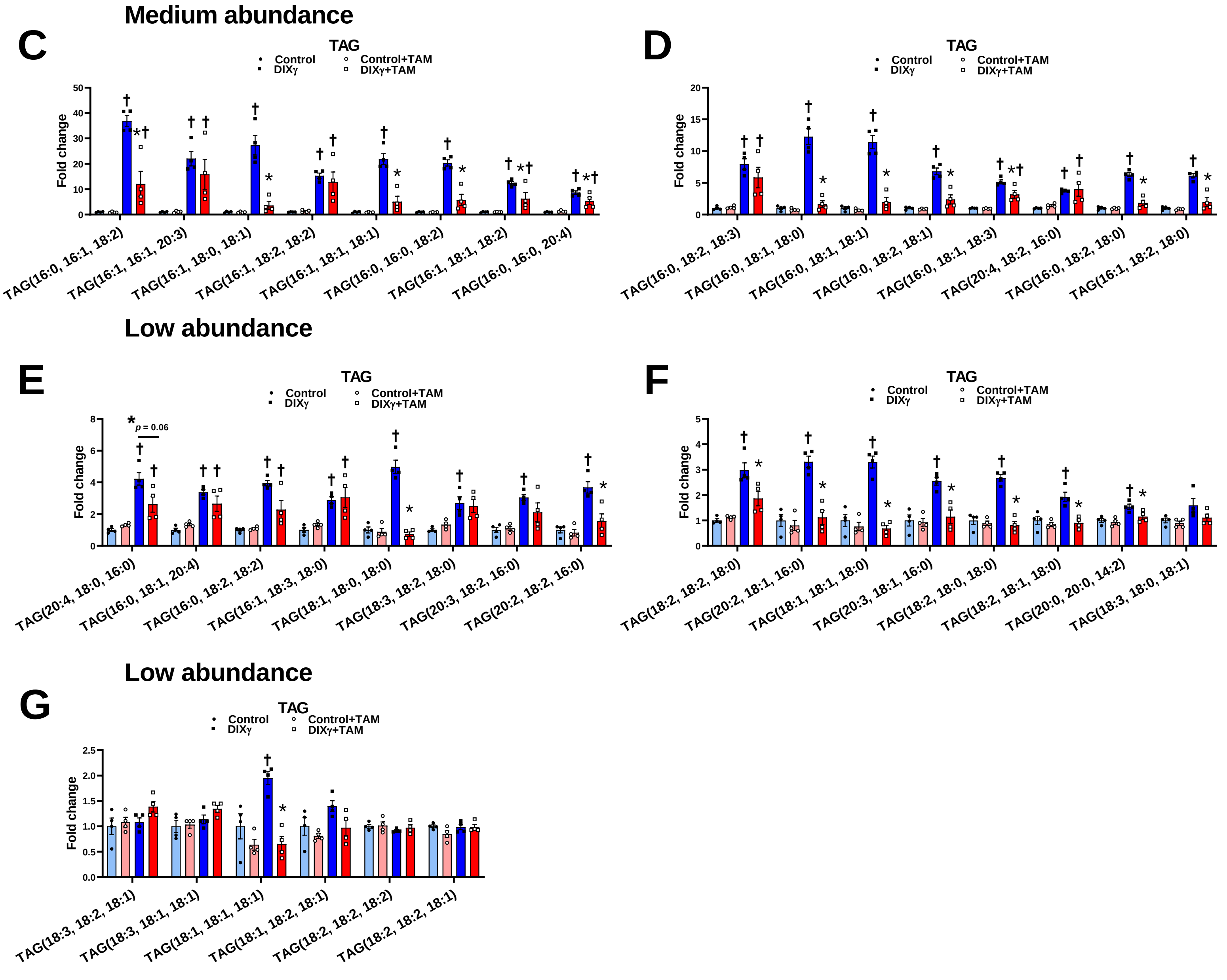


**Figure S4 continued: *Mogat1* knockout prevents accumulation of glycerolipids in differentiated SVF cells.** Stromal vascular fractions (SVF) were isolated from iWAT of *Mogat1* fl/fl mice. Confluent cells were treated with tamoxifen (4 nM) at the onset of differentiation (via DIXγ cocktail) and harvested after 10 days. **C-G:** TAG content was analyzed via LC-MS/MS and was lowered by *Mogat1* knockout. Data are expressed as mean ± S.E.M. * $p < 0.05$ vs. vehicle, † $p < 0.05$ *vs.* control undifferentiated cells; $n = 4$ biological replicates.

Figure S5

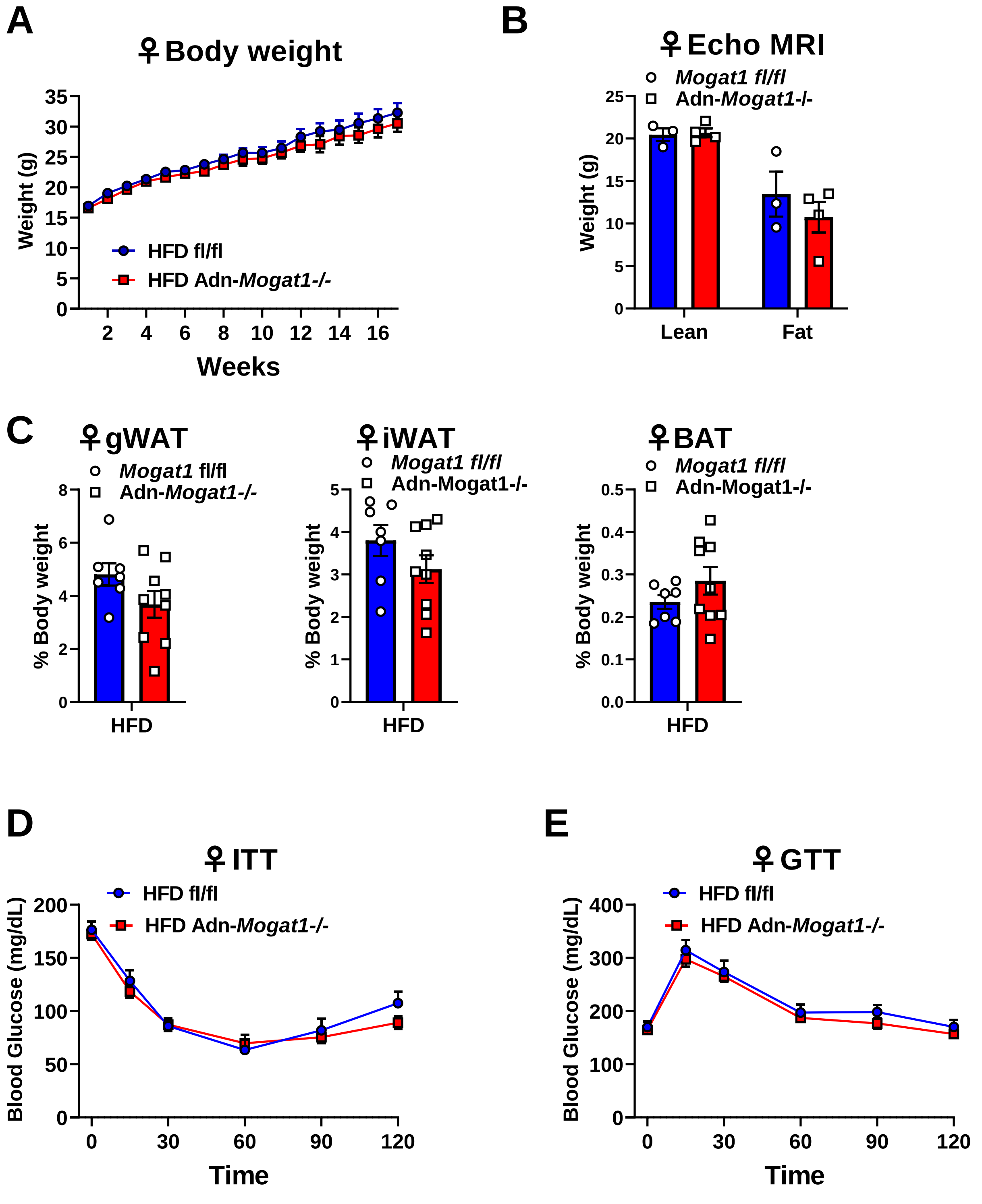


**Figure S5. Female Adipocyte-Specific *Mogat1* knockout mice gain similar weight on a long-term high fat diet (HFD) compared to littermate controls.** Female *Mogat1* fl/fl and littermate adiponectin Cre + (Adn-*Mogat1*-/-) mice were fed a 60% HFD starting at eight weeks of age. After 16 weeks of diet, mice were fasted for 4 hours prior to sacrifice and tissue collection. **A, B:** Weight gain and % lean or fat mass was unaffected by *Mogat1* knockout as measured by Echo MRI. **C:** gWAT and iWAT but not BAT were increased by HFD and are expressed as % total body weight. **D, E:** Insulin tolerance tests (ITT, 0.75 U/Kg lean mass, 4 hour fast) and glucose tolerance tests (GTT, 1 g/Kg lean mass, 5 hour fast) were similar between groups. Data are expressed as mean ± S.E.M. n = 7-9; n = 3-4 for Echo MRI.

Figure S6

A

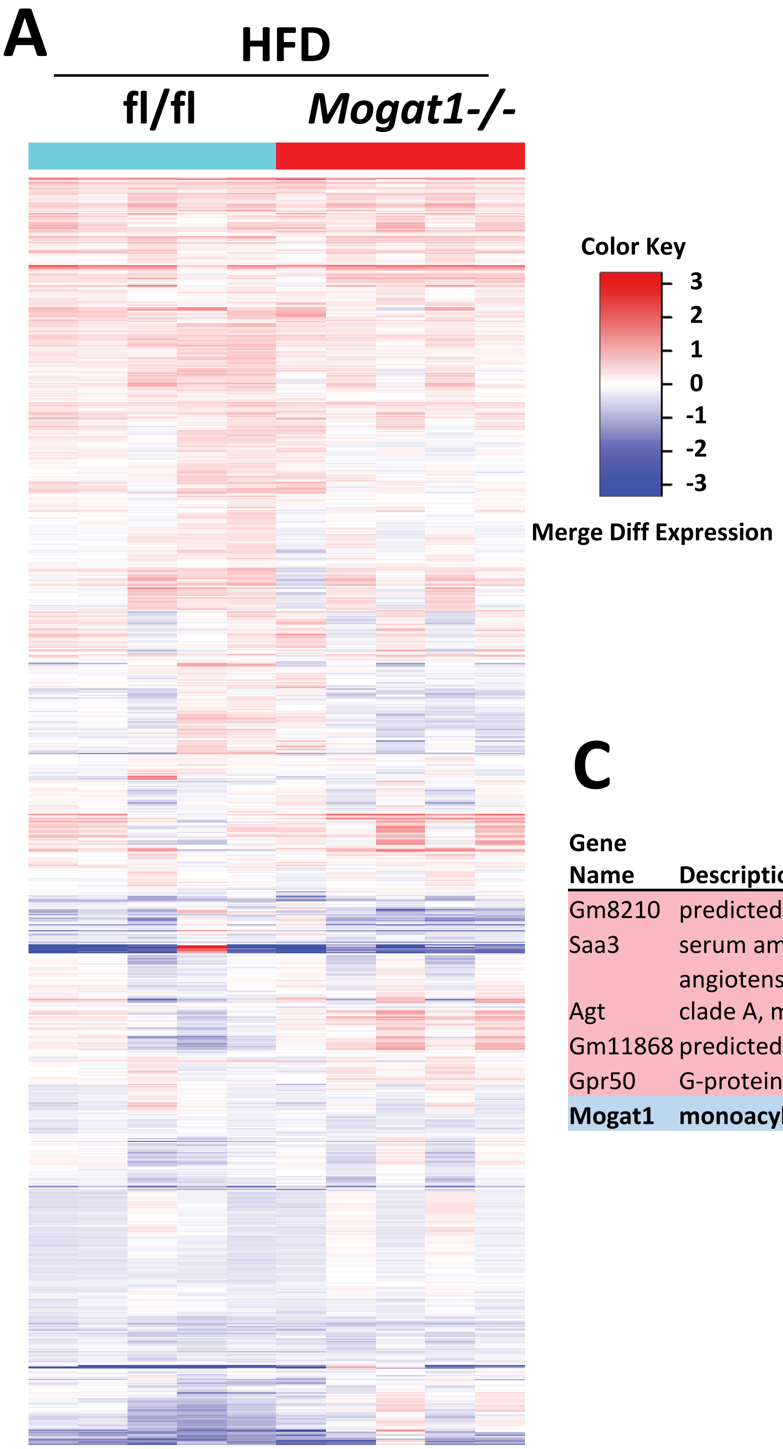


B

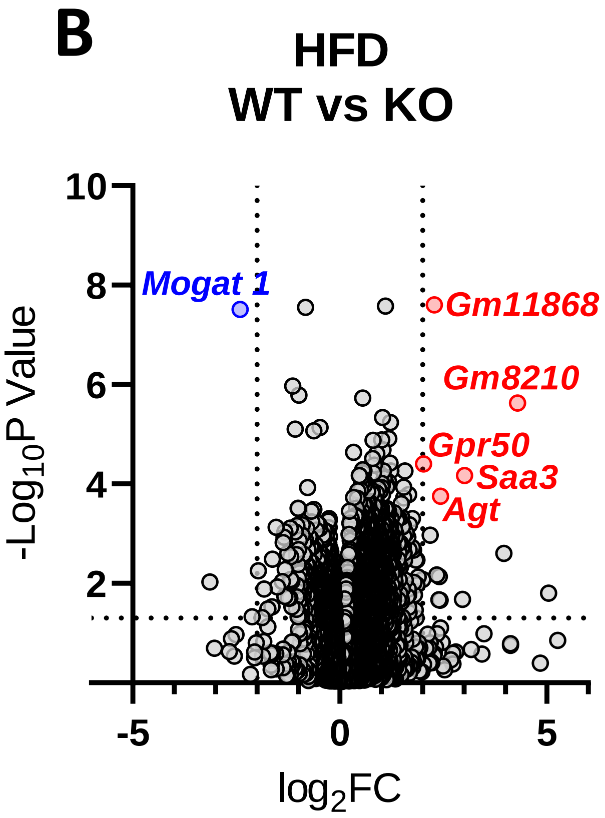


C

| Gene Name | Description | $Log_2FC$ | *p value* | *adj p* |
|---|---|---|---|---|
| Gm8210 | predicted pseudogene 8210 | 4.29 | 2.33E-06 | 4.50E-03 |
| Saa3 | serum amyloid A 3 | 3.02 | 6.82E-05 | 3.20E-02 |
| Agt | angiotensinogen (serpin peptidase inhibitor, clade A, member 8) | 2.43 | 1.78E-04 | 4.59E-02 |
| Gm11868 | predicted gene 11868 | 2.29 | 2.51E-08 | 1.19E-04 |
| Gpr50 | G-protein-coupled receptor 50 | 2.03 | 4.02E-05 | 2.82E-02 |
| **Mogat1** | **monoacylglycerol O-acyltransferase 1** | **-2.41** | 3.08E-08 | 1.19E-04 |

**Figure S6. Bulk RNA sequencing analysis of iWAT shows few differences between *Mogat1-/-* and littermate controls on a HFD. A:** Heatmap of merged differential expression data from iWAT from *Mogat1-/-* littermate controls on a HFD. **B:** Volcano plots of merged differential expression data was graphed as $\log_2$ fold change verses $-\log_{10}$ unadjusted *p* value. **C:** List of significantly changed genes. Gene expression changes were considered meaningful if they were > $\log_2$ fold change and had an adjusted *p* value of <0.05 vs. fl/fl mice on HFD.

Figure S7

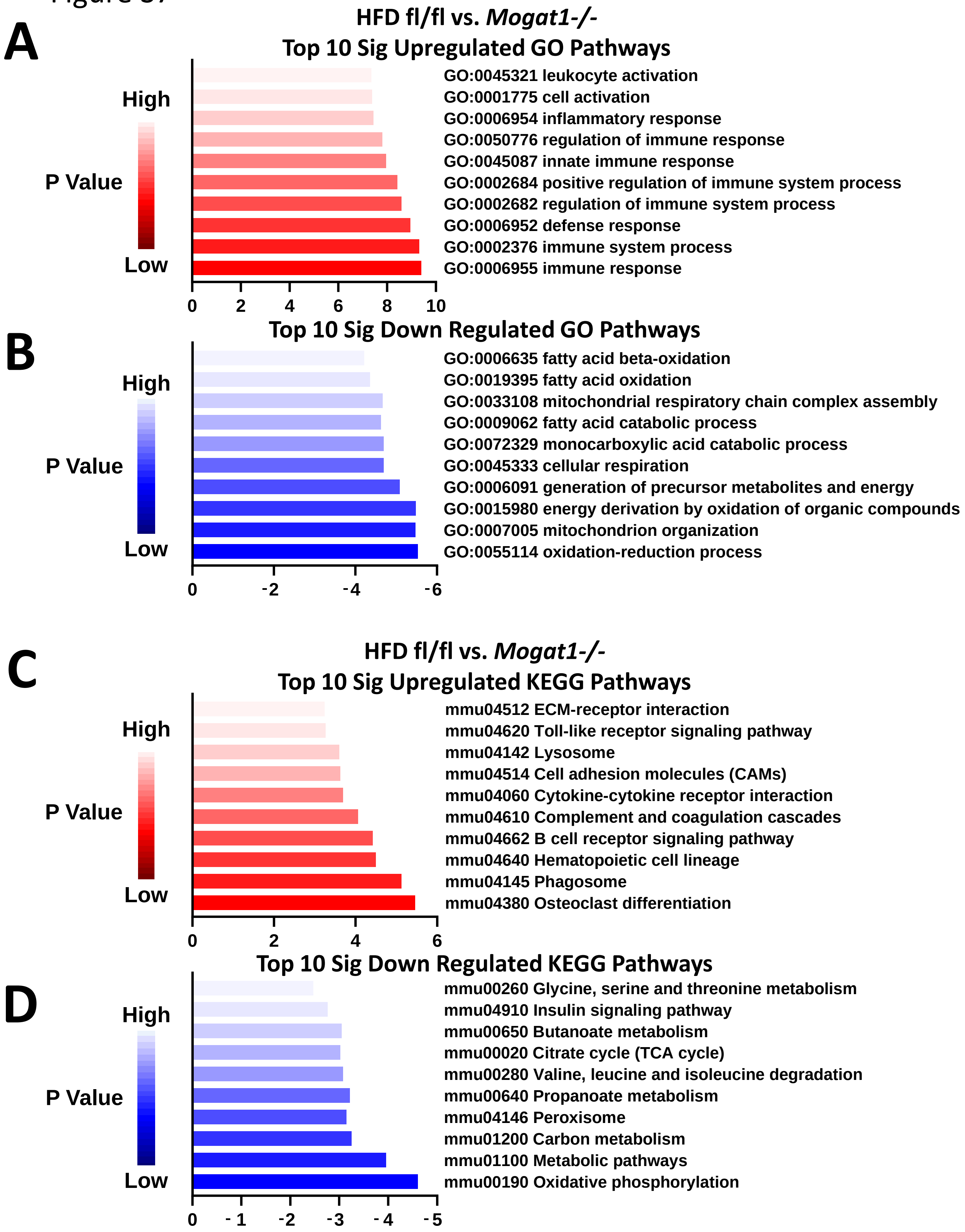


**Figure S7: *Mogat1* ablation in adipocytes alters metabolic pathways. A-D:** Graphical representation of pathway analysis showing top 10 significant signal direction changes of GO biological process and KEGG Signaling and Metabolism pathways. The color of the pathway label indicates the *p* value and the is the x-axis is the mean log fold change.
